# Adhesion-growth factor crosstalk regulates AURKB activation and ERK signaling in re-adherent fibroblasts

**DOI:** 10.1101/2021.04.28.441757

**Authors:** Siddhi Inchanalkar, Nagaraj Balasubramanian

## Abstract

Aurora kinases despite their similarity have distinct roles in the cell cycle, which is regulated by cell-matrix adhesion and growth factors. This study reveals loss of adhesion and re-adhesion to differentially regulate Aurora kinases. AURKB activation that drops on the loss of adhesion recovers on re-adhesion in serum-deprived conditions but not in the presence of serum growth factors. A rapid 30min serum treatment of serum-deprived cells blocks the adhesion-dependent recovery of AURKB, which negatively corelates with Erk activation. AZD mediated inhibition of AURKB in serum-deprived re-adherent cells promotes Erk activation and membrane ruffling, comparable to presence of serum. These studies thus define a novel adhesion-growth factor-dependent regulation of AURKB that controls adhesion-dependent Erk activation in re-adherent fibroblasts.

## Introduction

Aurora kinase A, B, and C belong to an evolutionarily conserved family of serine/threonine kinases. Aurora kinases although sharing high structure similarity, perform distinct functions owing to their distinct subcellular localization in cells. The mitotic kinases Aurora Kinase B (AURKB) and Aurora Kinase A (AURKA) share about 71 percent identity in their catalytic domain (Carmena and Earnshaw 2003). Despite this similarity, AURKB and AURKA are found to be differentially activated in response to a variety of upstream stimuli and also regulate distinct downstream pathways during mitosis (Barr and Gergely 2007; Goldenson and Crispino 2015; Lan *et al*. 2004; Nair *et al*. 2009). AURKB is involved in chromosome condensation, sister chromatid cohesion, mitotic spindle assembly, regulation of spindle assembly checkpoint, cytokinesis, and daughter cell spreading (Crosio *et al*. 2002; Ferreira *et al*. 2013). AURKA is essential in centrosome maturation, centrosome separation, the formation of bipolar spindle assembly, and G2-M transition (Terada *et al*. 2003). All these functions are crucial for cell proliferation, with de-regulation of these kinases leading to incomplete and improper cell division leading to aneuploidy and affecting cell viability (Khan *et al*. 2011; Umstead *et al*. 2017).

Integrin-dependent signalling is an important mediator of cell-cycle progression during the G1 phase, in particular (Assoian and Schwartz 2001; Bae *et al*. 2014; Jones *et al*. 2019; Klein *et al*. 2009; Moreno-Layseca and Streuli 2014). Extracellular forces can regulate cell-cycle checkpoints through a focal adhesion kinase (FAK)/Rac signalling module (Bae *et al*. 2014) and the Hippo pathway on changing matrix stiffness (Dupont *et al*. 2011) (Aragona *et al*. 2013). Integrin-mediated adhesion is further seen to regulate cytokinesis and the re-spreading and repulsive migration of daughter cells (Aszodi *et al*. 2003; Hognas *et al*. 2012; Mathew *et al*. 2014; Pellinen *et al*. 2008; Taneja *et al*. 2016). While β1 integrin influences cytokinesis, αVβ5-positive reticular adhesions are essential for the normal progression of mitosis. Patterned 2D and 3D matrices reveal local tissue shape and mechanics to regulate spatial differences in proliferation of cells (Aragona *et al*. 2013; Nelson *et al*. 2005; Nelson *et al*. 2006). Exogenous stretching of epithelial cells is seen to stimulate cell cycle progression from G1 to S (Benham-Pyle *et al*. 2015) and G2 to M (Gudipaty *et al*. 2017). Cells are thus able to sense their surrounding ECM environment through integrin-associated adhesion complexes and determine whether to proceed through S phase or enter mitosis. Growth factors that activate RTKs can regulate integrin-mediated cell adhesion, spreading, and migration (Klemke *et al*. 1994; Mainiero *et al*. 1996; Trusolino *et al*. 1998). Conversely, integrins also support the activation of growth factor signalling that regulates cell cycle progression in fibroblasts (Assoian and Schwartz 2001; Danen and Yamada 2001; Gu *et al*. 1992). Integrin mediated cell cycle progression is mediated by the regulation of cyclin D1 expression through multiple pathways involving Erk, PI-3K, and the Rho family GTPases Rac, cdc42, and Rho (Danen and Yamada 2001; Roovers *et al*. 1999) and the negative regulation of p21^cip1^ and p27^kip1^ (Bottazzi *et al*. 1999).

During mitosis, human cells undergo mitotic cell rounding, decreasing their adhesion to extracellular matrix substrates (Li and Burridge 2019; Uroz *et al*. 2018). Spatiotemporal regulation of various mitotic kinase activity aids in the extensive cytoskeletal remodelling mechanisms that prevent detachment from epithelia, while aiding successful completion of cytokinesis (Champion *et al*. 2017; Petridou *et al*. 2019). AURKB is a dynamic mitotic kinase that has been reported to have multiple roles during different phases of the cell cycle. Apart from its extensively studied functions as a part of chromosome passenger complex (CPC), in early G1 phase AURKB associated with the cell cortex regulates cell spreading. It interacts and phosphorylates a formin, FHOD1, that is known to be essential to organize cytoskeleton after cell division (Floyd *et al*. 2013). Ferreira et al, report that, in late cytokinesis, the presence of a gradient of AURKB activity ensures that microtubules (MT) in the furrow region remain phosphorylated and the ones in the vicinity of extracellular matrix stay de-phosphorylated, restricting the MT growth at the cell-matrix interphase. This allows for the stabilization of focal adhesion, aiding co-ordinated daughter cells spreading (Ferreira *et al*. 2013). A membrane raft protein Flotillin-1 has been recently shown to regulate AURKB activity and CPC function providing a direct point of regulation between extracellular cues and cell cycle progression (Gomez *et al*. 2010). It however remains unclear if and how integrin-mediated cell-matrix adhesion can regulate Aurora Kinase B activation, and if the presence of serum growth factors can affect this regulation. Also of interest would be determining if these effects are dependent on the cell cycle.

In this study, we have hence tested if and how cell-matrix adhesion regulates AURKA / AURKB activity in anchorage-dependent WT-MEFs and if this contributes to integrin-dependent signalling (ERK, FAK, AKT) and cell spreading. The role serum growth factors have in regulating adhesion-dependent AURKB activation and the possible contribution cell cycle profile has on this regulation were also evaluated.

## Materials and Methods

### Reagents

Human plasma fibronectin (Cat#F2006), nocodazole (Cat # M1404), DMSO (Cat # D2438), DAPI (Cat# D9542), Propidium iodide (Cat# P4170), and sparfloxacin (Cat# 56968) were purchased from Sigma, and Phalloidin-Alexa-488 (Cat# A12379) was from Molecular Probes (Invitrogen). Fluoromount-G used to mount cells for imaging was obtained from Southern Biotech (Cat# 0100-01). AZD1152 (AURKB inhibitor Cat# SML0268) was purchased from Sigma. RNAase-A (Cat# 9001-99-4) was purchased from USB corporation. Immobilon Western blot substrate (Cat# WBKLS0500) was purchased from Millipore. Antibodies used for western blotting including anti-phospho-aurora A (Thr288)/aurora B (Thr232)/ aurora C (Thr198) (Cat #2914), anti-AURKB (Cat #3094), anti-phospho AKT-Ser473 (Cat# 9271) (1:2000 dilution), anti-phospho-FAK-Tyr397 (Cat# 3283), anti-phospho-p44/p42 ERK1/2 (Thr202/Tyr204) (Cat# 4370) (1:2000 dilution), anti-p44/p42 ERK1/2 (Cat# 4695) (1:2000 dilution), anti-FAK (Cat#3285) and anti-AKT (Cat# 4691) were purchased from Cell Signaling Technologies and used at 1:1000 dilution unless mentioned otherwise. Anti-AURKA (Cat #610939) was purchased from BD Transduction Laboratories used at 1:1000 dilution. Anti-beta actin (Cat# Ab3280) antibody was purchased from Abcam used at 1:2000 dilution. Secondary antibodies conjugated with HRP were purchased from Jackson Immunoresearch and were used at a dilution of 1:10000. Antibody used for immunofluorescence include anti-phospho-p44/p42 ERK1/2 (Thr202/Tyr204) (Cat# 4370) from Cell signaling technologies used at 1:200 dilution. Secondary antibodies with Alexa conjugate (488 or 594) were purchased from Invitrogen Molecular Probes (Cat. No. # A12379 and A12381) and were used at a dilution of 1:1000.

### Cell Culture

Wild-type Mouse embryonic fibroblasts (WT-MEFs) obtained from Dr. Richard Anderson (University of Texas Health Sciences Centre, Dallas TX), were cultured in DMEM (Cat# 11995-065) supplemented with 5% (v/v) fetal bovine serum (FBS) (Cat# 26140-079) and 1% (v/v) penicillin-streptomycin (Cat# 15140-122) (all from Invitrogen) at 37°C under 5% CO_2_ humidified atmosphere.

### Suspension and replating of WTMEFs in serum starved or with serum conditions

For suspension assay, serum-starved (0.2% FBS) WT-MEFs (for at least 12 hours) were detached with 1X trypsin-EDTA diluted with low serum medium and held in suspension for 30 minutes with 1% methylcellulose in low serum DMEM. Post incubation for respective time points cells were carefully washed twice with 0.2% FBS DMEM at 4°C to avoid clumping and collected. These washed cells were re-plated on dishes or coverslips coated overnight with fibronectin at 4°C (2μg/ml or 10μg/ml as indicated in figure legends) for 15 minutes or 4 hours. Cells were similarly grown and processed with serum (10% FBS) when required. When needed for western blotting these cells were lysed in the required volume of 1X laemmli, heated at 95°C for 5mins and stored at -80°C. For confocal microscopy cells were fixed with 3.5% paraformaldehyde (PFA) for 15 mins at room temperature (RT), washed with PBS thrice, stained, and mounted.

### Serum stimulation suspension assay

For serum stimulation suspension assay, serum-starved (0.2% FBS) WT-MEFs (for at least 12 hours) were detached with 1X trypsin-EDTA diluted with low serum medium and held in suspension for 30 minutes with 1% methylcellulose in low serum DMEM. Post 30mins of suspension, a predetermined amount of 100% FBS was added to the suspension culture to adjust the final FBS concentration to 10%. Cells were suspended in 10% complete FBS or 10% heat-inactivated FBS (heat inactivation was done at 56°C for 30mins before the start of the experiment) media for additional 15mins to bring the total suspension timepoint to 45 mins. Post 45mins (30mins low serum + 15mins 10% serum) incubation for respective time points cells were carefully washed twice with 10% FBS DMEM at 4°C to avoid clumping and collected. These washed cells were re-plated on dishes coated overnight with fibronectin at 4°C (10μg/ml) for 15minutes or 4 hours. When needed for western blotting these cells were lysed in the required volume of 1X laemmli, heated at 95°C for 5mins, and stored at -80°C.

### Cell cycle analysis by flow cytometry

For evaluating adhesion-dependent cell cycle profile, WT-MEFs grown in either 0.2% FBS DMEM or 10% FBS DMEM were detached using trypsin and held in suspension for 30 minutes. Post suspension cells were carefully washed, collected, and divided into three equal proportions: one processed immediately as suspension time point (SUS), one re-plated on 10μg/ml fibronectin-coated dishes for 15minutes (FN 15’), and last re-plated on 10μg/ml fibronectin-coated dishes for 4 hours (SA). These cells were washed twice with PBS and fixed using chilled 70% ethanol and stored at 4°C till further use (not more than 18hours). On the day of flow cytometer measurement cells were treated with 100μg/ml RNAse A and labeled with 10μg/ml of propidium iodide. DNA content was analyzed for cell cycle status in BD LSRFortessa SORP cell analyzer. 10000 events were recorded for each treatment and time point to obtain the percentage of cells in different phases of the cell cycle. The cell cycle profiles were calculated by using ModFit software that gives percentages of cells in G0-G1, S, and G2-M phase and were compared across different treatments and time points. This method was used to evaluate the cell cycle profile of WT-MEFs that are stable adherent, held in suspension for 30 minutes, and re-adherent on fibronectin for 15 minutes in the presence and absence of serum growth factors.

### Western Blot Detection of Proteins

30 μl of lysate was resolved by 12.5% SDS-PAGE and transferred to PVDF membrane (Millipore). Blots were blocked with 5% milk in 0.1% Tween-20 containing Tris-buffered saline (TBST) for 1 hour at room temperature and incubated with required antibody diluted in 5% BSA at 4°C overnight. Blots were then washed thrice with TBST and incubated with anti-mouse HRP (AURKA) and anti-rabbit HRP (AURKB, Phospho-AURK A/B/C, ERK, FAK, Phospho-ERK, Phospho-FAK) for an hour, followed by detection using chemiluminescent substrates from Millipore. LAS4000 (Fujufilm-GE) was used to image the blots and densitrometric band analysis was done using Image-J software (NIH). Prism Graphpad software was then used to do the statistical analysis.

### Immunofluorescence assay

WT-MEFs were serum-starved with 0.2% FBS containing DMEM (low serum DMEM) for 12 hours, detached, held in suspension for 30mins in the absence or presence of AZD1152, and re-plated on fibronectin (2μg/ml) for 15mins. Re-adherent cells were fixed with 3.5% paraformaldehyde after 15minutes of re-plating. Cells were permeabilized with PBS containing 5% BSA and 0.05% Triton-X-100 for 15 minutes and blocked with 5% BSA for 1 hour at room temperature followed by incubation with 1:200 rabbit anti-phospho-p44/p42 ERK1/2 (Thr202/Tyr204) antibody in 5% BSA for 3 hours. Cells were finally stained with 1:1000 diluted secondary antibodies (anti-mouse Alexa-568) and 1:500 diluted phalloidin-Alexa 488 for 1 hour at room temperature. All incubations were done in a humidified chamber. Washes were done with 1X PBS at room temperature. Stained and washed coverslips were mounted with Fluoromount-G (Southern Biotech) and imaged using a Zeiss LSM 710 laser confocal-Anisotropy or LSM780 multiphoton microscope with a 63x objective.

### Statistical Analysis

All the analysis was done using Prism Graphpad analysis software. Statistical analysis of data was done using the two-tailed unpaired Student’s T-test for non-normalized data and when normalized to respective controls using the two-tailed single sample T-test. Distribution profile data was analysed using Chi-square test.

## Results

### Cell-matrix adhesion differentially regulates AURKB vs AURKA activation

To evaluate if cell-matrix adhesion can regulate Aurora Kinase A (AURKA) and Aurora Kinase B (AURKB) activity, serum-starved stable adherent (SA) wild-type mouse embryonic fibroblasts (WT-MEFs) were detached and held in suspension for 30mins (with 1% methylcellulose) (SUS 30’), and re-plated on fibronectin for 15mins (FN 15’). In the absence of serum growth factors, re-adhesion of cells is known to activate integrin-dependent downstream signalling. The effect that loss of adhesion and re-adhesion has on Aurora Kinase activity was tested by evaluating their phosphorylation at the Threonine 288 (AURKA) and Threonine 232 (AURKB) residues, known to be necessary for their activation (Ohashi *et al*. 2006; Yasui *et al*. 2004). In serum-starved WT-MEFs, AURKA activity drops (∼60%) upon loss of adhesion and does not recover on re-adhesion (Figure 1A), on the other hand, AURKB activity drops (∼40%) upon loss of adhesion and is restored upon re-adhesion to fibronectin (Figure 1B). AKT activation known to be regulated by adhesion expectedly drops on the loss of adhesion (SUS 30’) and recovers on re-plating for 15mins (FN 15’) (Figure S1A). This suggests that integrin-dependent adhesion differentially regulates AURKA *vs* AURKB activity in WT-MEFs.

**Figure 1.**
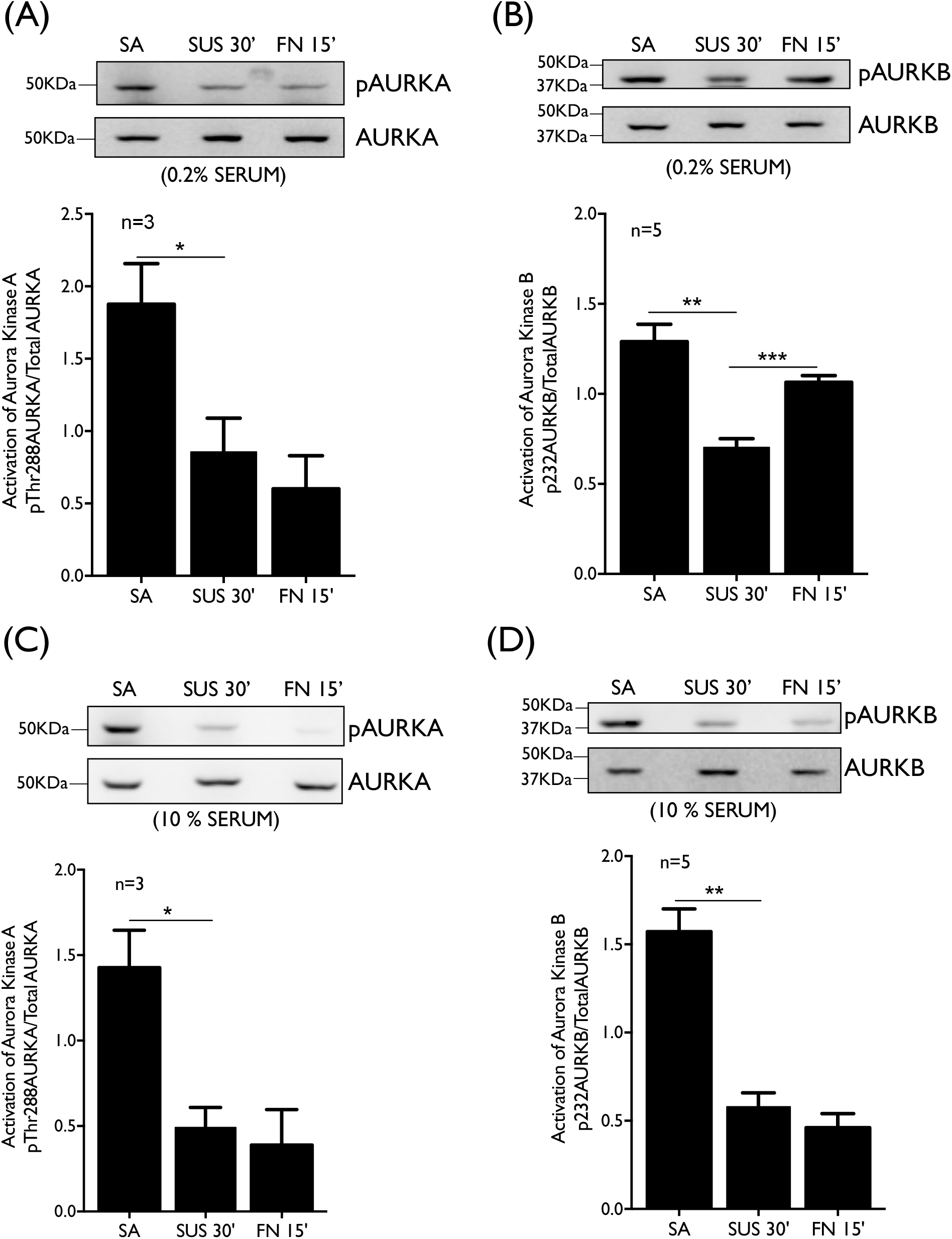
Adhesion-growth factor crosstalk dependent regulation of AURKA and AURKB. Western blot detection (upper panel) and quantitation (lower panel) of (A and C) phosphorylation on Threonine 288 residues of AURKA (pAURKA), total AURKA and, (B and D) phosphorylation on the Threonine 232 residues of AURKB (pAURKB), total AURKB in the lysates from WT-MEFs stable adherent (SA), suspended for 30 mins (SUS 30’) and re-adherent on fibronectin for 15mins (FN 15’) in presence of 0.2% (A and B) and 10% (C and D) FBS DMEM. The ratios of pAURKA/AURKA and pAURKB/AURKB are represented in the graph as mean ± SE from at least three independent experiments. Statistical analysis of all the above data was done using a two-tailed paired Student’s t-test and significance if any was represented in the graph (* p-value <0.05, ** p-value <0.01, *** p-value <0.0001).

### Adhesion-growth factor crosstalk regulates Aurora Kinase B activity

Knowing the overlap between adhesion and growth factor-mediated signaling, we tested the role serum growth factors could have on the adhesion-dependent regulation of Aurora kinases. On the loss of adhesion (suspension 30min) activation of both AURKA and AURKB, seen to drop in serum-deprived conditions, (Figure 1A and 1B) continue to decrease in the presence of serum growth factors (Figure 1C and 1D). AURKA activity that does not recover on re-adhesion to fibronectin for 15mins, in the absence of serum, is also unaffected by serum growth factors (Figure 1A and 1C). However, the recovery of AURKB activity on re-adhesion in serum-deprived conditions (Figure 1B) is interestingly lost in the presence of serum growth factors (Figure 1D). The AURKB activity in suspended cells without and with serum was comparable (Figure S1C) and could hence not affect their recovery on re-adhesion. The drop in AKT activity on the loss of adhesion is also reduced in the presence of serum, while its recovery on re-adhesion is retained (Figure S1B). Together they suggest integrin-mediated adhesion-dependent regulation of AURKB to indeed be affected by serum growth factors.

### Role of the cell cycle in adhesion-growth factor-mediated AURKB activation

Knowing the role of the cell cycle in regulation of Aurora kinases and the impact adhesion and growth factors could have on the same, we asked if and how their changing cell cycle profile could affect the observed regulatory crosstalk. Serum starvation of WT-MEFs, as expected, synchronize cells at the G1-S phase (Campisi *et al*. 1984; Griffin 1976; Langan and Chou 2011; Langan *et al*. 2017) with 76 ± 2.3 % of stable adherent cells in the G1-G0 phase and 3 ± 0.8 % in the G2-M phase (Figure 2A). This profile does not change significantly when the cells are detached and held in suspension for 30 mins, followed by re-plating on fibronectin for 15mins (Figure 2A). The absence of any significant change in the cell cycle profile of fibroblasts suggests the regulation of Aurora Kinase B by adhesion is indeed independent of the cell cycle.

**Figure 2:**
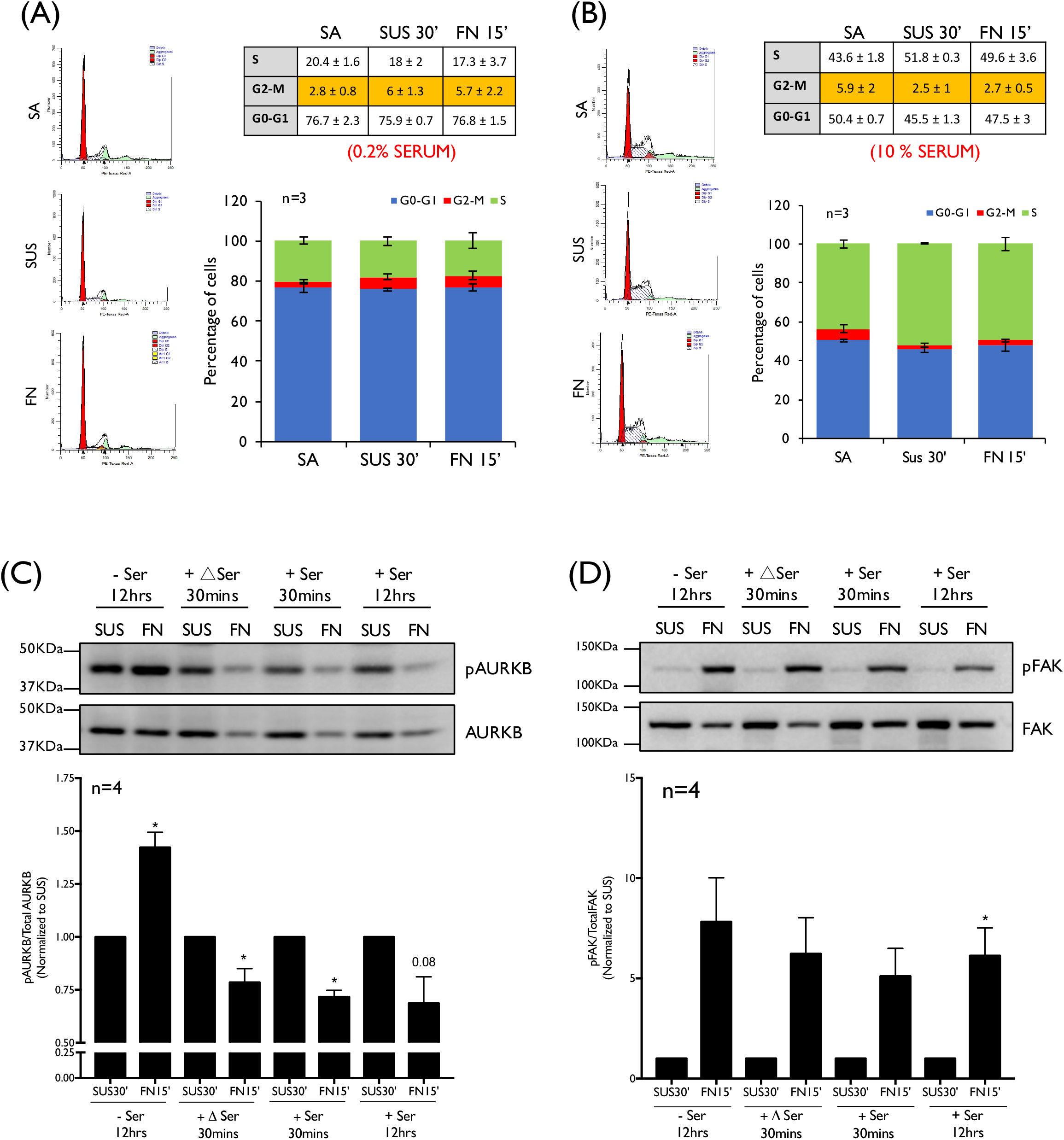
Adhesion-growth factor dependent regulation of Cell cycle profile and AURKB activation. Representative histogram of 3 independent experiments and percentage of cells present in G2-M, S and G0-G1 phase are shown in table and graph from stable adherent (SA), suspended for 30 mins (SUS 30’) and re-adherent on fibronectin for 15mins (FN 15’) in (A) serum starved and (B) with serum WT-MEFs. The graph represents the mean ± SE from at least 3 independent experiments. Statistical analysis was done using a two-tailed unpaired Student’s t-test and significance represented (* p-value <0.05, ** p-value <0.01). (C and D) Western blot detection (upper panel) and quantitation (lower panel) of (C) phosphorylation on the Threonine 232 residues of AURKB (pAURKB), total AURKB and, (D) phosphorylation on Tyrosine 397 residues of FAK (pFAK) and total FAK in the lysates from: serum starved WT-MEFs (-Ser 12hrs) suspended for 45mins and re-plated on fibronectin in presence of 0.2% serum, serum starved WT-MEFs suspended for 30 mins followed by 15mins in heat inactivated 10% FBS DMEM (+ΔSer 30mins) and re-adherent on fibronectin for 15mins (FN 15’) in presence of heat inactivated 10% serum growth factors, serum starved WT-MEFs suspended for 30 mins followed by 15mins in 10% FBS DMEM (+Ser 30mins) and re-adherent on fibronectin for 15mins (FN 15’) in presence of 10% serum growth factors or WT-MEFS suspended and replated in presence of 10% FBS DMEM. The ratios of pAURKB/AURKB and pFAK/FAK were normalized to respective SUS (equated to 1), and these values are represented in the graph as mean ± SE from four independent experiments. Statistical analysis of all the above data was done using the single sample t-test for normalized data and two-tailed unpaired Student’s t-test for non-normalized data and significance if any was represented in graph (* p-value <0.05).

To further establish the differential regulation of AURKB we tested adhesion-dependent regulation of the cell cycle profile of WT-MEFs in the presence of serum. 50 ± 0.7 % of stable adherent WT-MEFs were seen to be in the G1-G0 phase and 43 ± 1.8 % in the S phase (Figure 2B). This profile did not change significantly as these cells were held in suspension for 30 mins, followed by re-plating on fibronectin for 15mins (Figure 2B). In the presence of serum (Figure 2B), the cell cycle profile was expectedly different as compared to that seen in the absence of serum (Figure 2A).

The expression levels and activation of Aurora Kinases are seen to vary at different phases of the cell cycle (Carmena and Earnshaw 2003; Goldenson and Crispino 2015). A small but significant change in the cell cycle profile that the presence of serum causes could in part contribute to the differential effect serum growth factors have on adhesion-dependent AURKB activation. To evaluate this possibility, we tested if rapid (15min) stimulation of serum-deprived cells with serum growth factors, unlikely to affect their cell cycle profile, can affect the re-adhesion mediated regulation of AURKB. Serum-deprived (0.2% FBS DMEM) WT-MEFs suspended for 30mins were treated for 15min with 10% FBS and re-plated with 10% FBS DMEM on fibronectin for 15min. This treatment we find was enough to prevent the recovery of AURKB activity upon re-adhesion (Figure 2C) as seen in WT-MEFs grown with 10% serum (Figure 2C and 1D). This could have implications for serum growth factor-mediated fibroblast function. We hence tested if heat inactivation of serum (56°C, 30mins) can affect the adhesion-dependent AURKB activation (+ΔSer 30mins) and find that it does not (Figure 2C). The AURKB activity in suspended cells without and with serum and on serum stimulation was comparable (Figure S2A) and could hence not contribute to the recovery observed on re-adhesion. Focal adhesion kinase (FAK) activity is known to be regulated by integrin-mediated adhesion (Eliceiri 2001; Oktay *et al*. 1999) and was expectedly found to drop upon loss of adhesion comparably (Figure S2B) and recover back upon re-plating on fibronectin irrespective of presence or absence of serum growth factors or heat-inactivated serum (Figure 2D). This confirms serum growth factors to inhibit adhesion-dependent activation of AURKB, independent of the cell cycle.

### Role of AURKB in regulating adhesion-growth factor-dependent ERK activation

To evaluate the functional relevance adhesion and growth factor-mediated regulation of AURKB activation could have in cells, we investigated if and how it could affect adhesion-dependent signaling or function. We tested the effect absence and presence of serum has on re-adhesion mediated activation of ERK and FAK (Eliceiri 2001) in WT-MEFs and the role AURKB activation could have in mediating the same. In the presence and absence of serum, FAK activation drops on the loss of adhesion comparably (Figure S3C) and recovers on re-adhesion (Figure 3C and 2D). Adhesion-dependent ERK activation in WT-MEFs was however seen to be differentially affected by the presence or absence of serum (Figure 3A), like AURKB (Figure 3B, 1B, and 1D). In low serum conditions ERK activation that drops on the loss of adhesion, further decreases on re-adhesion to fibronectin (Figure 3A) as AURKB activation increases (Figure 3B). Basal ERK activity was found to be significantly higher in WTMEFs in the presence of serum (Figure 32A) but shows no significant change in its activation on the loss of adhesion or re-adhesion (Figure 3A). In contrast, AURKB activation is seen to be drop significantly in suspended cells and stays low on re-adhesion (Figure 3B) in the presence of serum. This suggests the presence of a distinct inverse correlation between ERK and AURKB activation in re-adherent cells.

**Figure 3:**
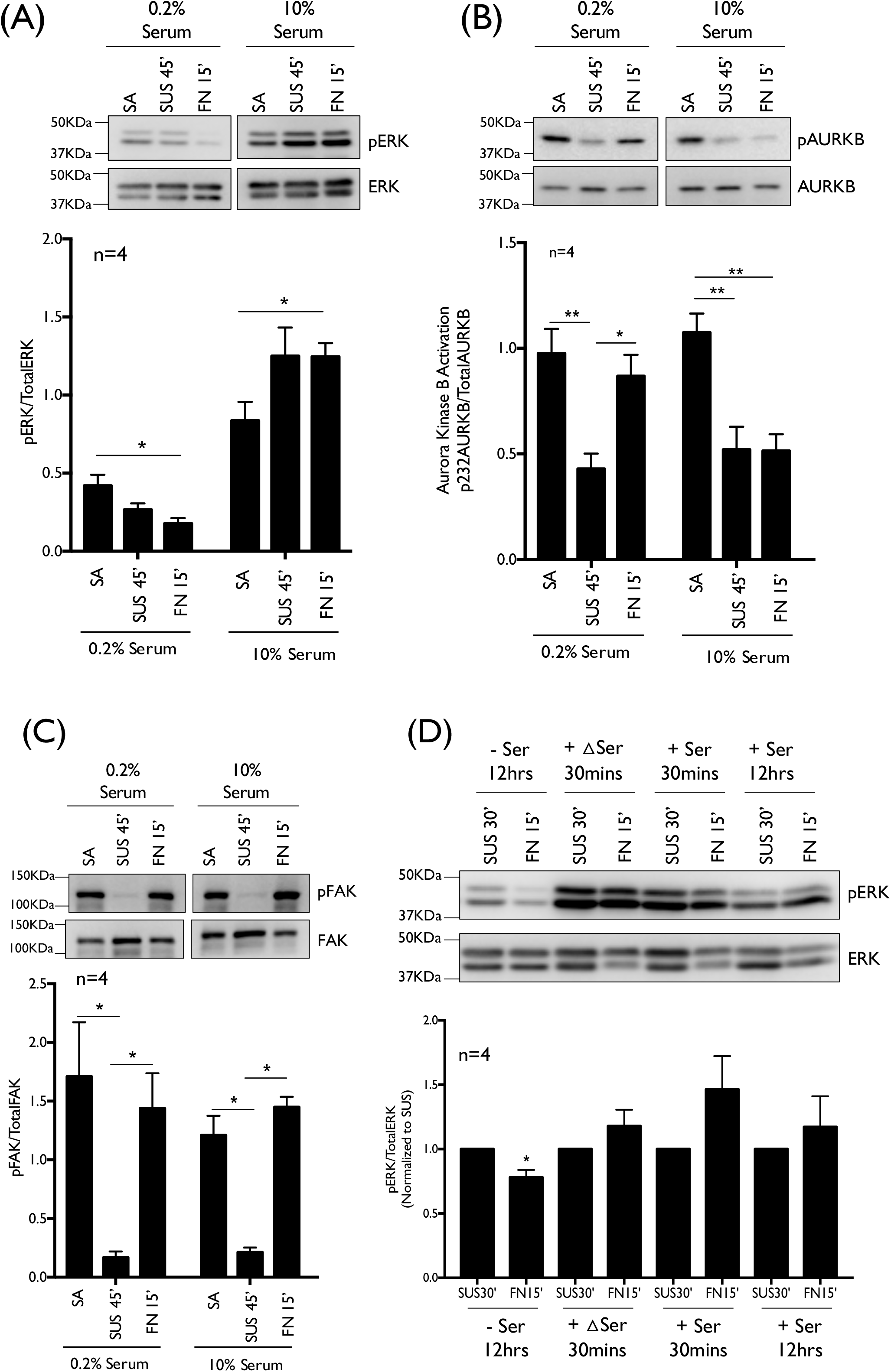
Adhesion-growth factor dependent regulation of ERK activity. Western blot detection (upper panel) and quantitation (lower panel) of (A) phosphorylation on Threonine202/Tyrosine204 residues of p44/p42 ERK1/2 (pERK) and Total p44/p42 ERK1/2 (TotalERK), (B) phosphorylation on the Threonine 232 residues of AURKB (pAURKB), total AURKB and, (C) phosphorylation on Tyrosine 397 residues of FAK (pFAK) and total FAK in the lysates from serum-starved WT-MEFs suspended for 45mins and re-plated on fibronectin in presence of 0.2% serum and WT-MEFS suspended and re-plated in presence of 10% FBS DMEM. (D) Western blot detection (upper panel) and quantitation (lower panel) of phosphorylation on Threonine202/Tyrosine204 residues of p44/p42 ERK1/2 (pERK) and Total p44/p42 ERK1/2 (TotalERK), in the lysates from, serum starved WT-MEFs (-Ser 12hrs) suspended for 45mins and re-plated on fibronectin in presence of 0.2% serum, serum starved WT-MEFs suspended for 30 mins followed by 15mins in heat inactivated 10% FBS DMEM (+ΔSer 30mins) and re-adherent on fibronectin for 15mins (FN 15’) in presence of heat inactivated 10% serum growth factors, serum starved WT-MEFs suspended for 30 mins followed by 15mins in 10% FBS DMEM (+Ser 30mins) and re-adherent on fibronectin for 15mins (FN 15’) in presence of 10% serum growth factors or WT-MEFS suspended and re-plated in presence of 10% FBS DMEM The ratios of pERK/TotalERK, pAURKB/AURKB, and pFAK/FAK are represented in the graph as mean ± SE from four independent experiments. Statistical analysis of all the above data was done using the single sample t-test for normalized data and two-tailed unpaired Student’s t-test for non-normalized data and significance if any was represented in graph (* p-value <0.05, ** p-value<0.01).

To confirm this, we evaluated the regulation of ERK on stimulation of serum-deprived cells with serum as well as heat-inactivated serum for 15minutes. Both cause a drop in AURKB activity on re-plating as compared to serum-starved cells (Figure 2C). This regulation of AURKB is comparable to that seen in cells grown with serum (Figure 3B). ERK activity in re-adherent cells is promoted in the presence of serum (Figure 3D) further suggesting the adhesion-dependent AURKB activation to inversely regulates ERK activity in re-adherent cells. To further confirm the adhesion-dependent AURKB-ERK crosstalk, we tested the effect inhibition of AURKB activity has on ERK activation. We used a specific AURKB small-molecule inhibitor AZD1152 (Wilkinson *et al*. 2007) to treat serum-deprived re-adherent cells (where AURKB is activated) (Figure 1B) and compared their ERK activation to serum treated cells. AZD1152 treatment for 30mins inhibits AURKB (Figure 4A and S4B), which causes a significant increase in ERK activity upon re-adhesion, comparable to serum treated cells (Figure 4B). No significant effect of AURKB inhibition on ERK activation was seen in suspended cells (Figure S4C) or on FAK activation in suspended as well as re-adherent WT-MEFs (Figure S4A and S4D). Taken together, this confirms the presence of a unique AURKB-ERK regulatory crosstalk in re-adherent cells.

**Figure 4:**
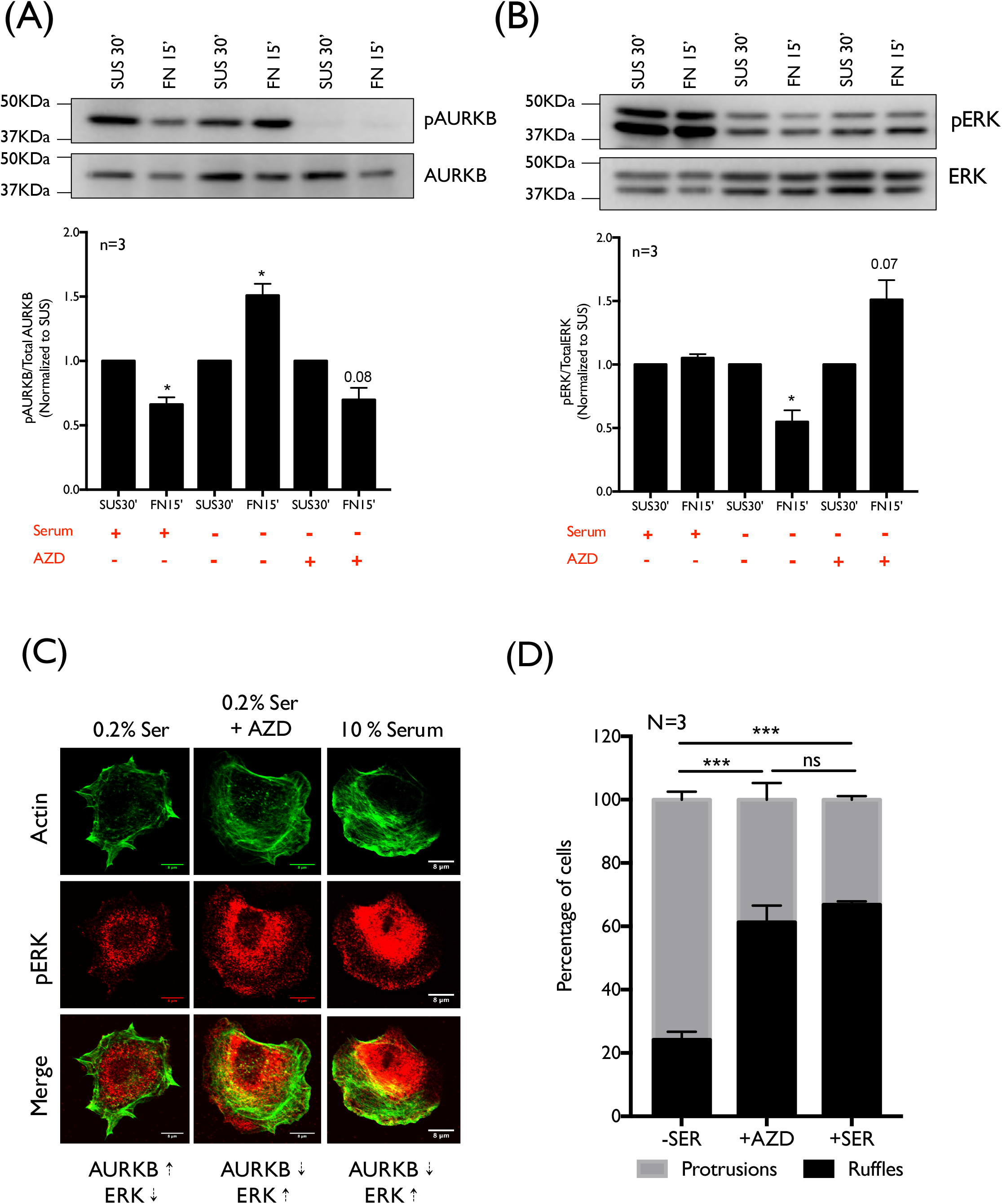
Effect of AURKB inhibition on re-adherent ERK activity, localization, and membrane ruffling in WT-MEFs. Western blot detection (upper panel) and quantitation (lower panel) of (A) phosphorylation on the Threonine 232 residues of AURKB (pAURKB), total AURKB and, (B) phosphorylation on Threonine202/Tyrosine204 residues of p44/p42 ERK1/2 (pERK) and Total p44/p42 ERK1/2 (TotalERK) in the lysates from serum-starved WT-MEFs suspended for 30mins (with or without 2μM AZD1152) and re-plated on fibronectin in presence of 0.2% serum (with or without 2μM AZD1152) and WT-MEFS suspended and replated in presence of 10% FBS DMEM. The ratios of pERK/TotalERK and pAURKB/AURKB are represented in the graph as mean ± SE from three independent experiments. Statistical analysis of all the above data was done using the single sample *t*-test and significance if any was represented in graph (* p-value <0.05). (C) pERK detected using specific antibody against phosphorylation on Threonine202/Tyrosine204 residues of p44/p42 ERK1/2 (pERK) at membrane ruffles in re-adherent spreading WT-MEFs in presence of 0.2% FBS, 0.2% FBS + 2 μM AZD1152 or 10% FBS DMEM. (D) Percentage distribution of cells with ruffled and protrusion phenotypes in re-adherent WT-MEFs in presence of 0.2% FBS or 0.2% FBS + 2 μM AZD1152, or 10% FBS DMEM, was determined by counting 100 cells per time point from three independent experiments. Statistical analysis was done using Chi-square test two-tailed for distribution profile (D) data and p-values are as indicated (**p<0.01, *** p < 0.001). Scale bar in (C) is set at 8 μm.

We further evaluated the localization of active phosphorylated ERK (pERK-Thr202/Tyr204) in re-adherent cells and tested the impact AZD1152 mediated AURKB inhibition and the resulting stimulation of ERK activation has on the same. Serum-deprived re-adherent cells make significantly fewer membrane ruffles unlike serum-treated WTMEFs (Figure 4C and 4D). Active pERK is seen to prominently localize at membrane ruffles in serum treated cells where AURKB activation is blocked. In serum-deprived AZD1152 treated re-adherent cells, the inhibition of AURKB now causes cells to have a distinct ruffling phenotype with pERK actively recruited to membrane ruffles (Figure 4C and 4D). Together these results confirm adhesion-dependent AURKB-ERK regulatory crosstalk to control ERK activity and localization downstream of integrin-mediated adhesion (Figure 5).

**Figure 5.**
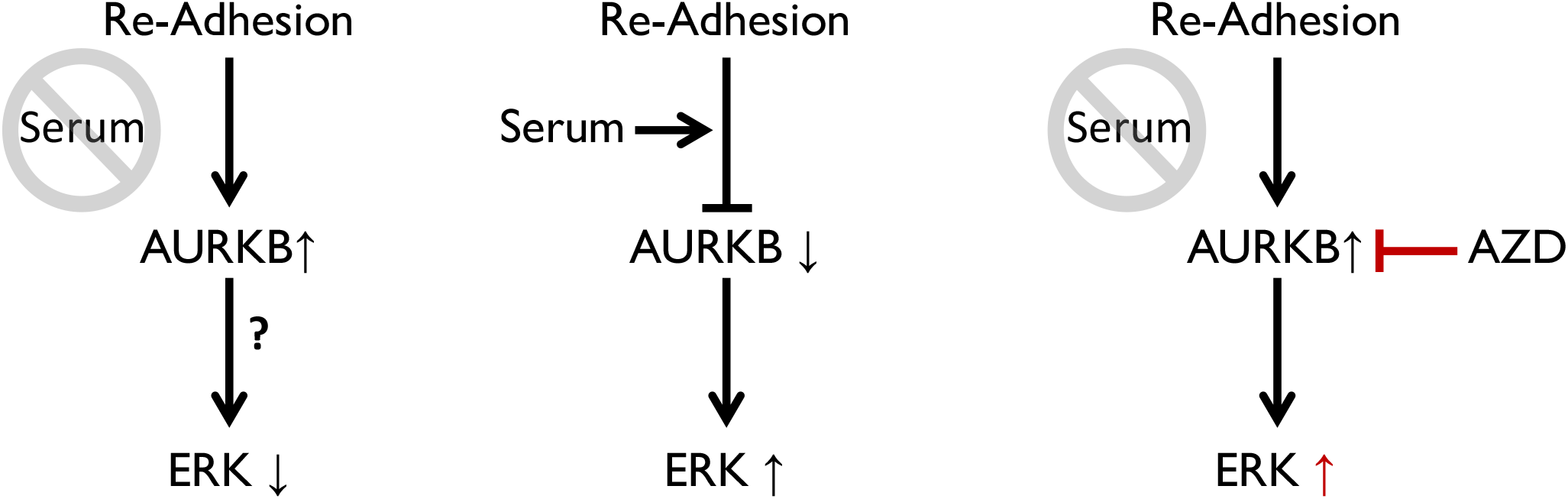
Adhesion and serum growth factor dependent regulation of AURKB and ERK in re-adherent cell. This schematic captures how re-adhesion to fibronectin in the absence of serum triggers the activation of AURKB to reciprocally suppress Erk activation. This is reversed by the presence of serum that prevents re-adhesion mediated recovery of AURKB activation increasing Erk activation. AZD mediated inhibition of AURKB activation in serum deprived re-adherent cells promotes Erk activation, confirming this regulatory crosstalk.

## Discussion

Aurora kinases are important regulators of cell division, single and collective cell migration (Barr and Gergely 2007; Carmena and Earnshaw 2003; Goldenson and Crispino 2015; Mahankali *et al*. 2015; Zhu *et al*. 2014) all of which are known to also be influenced by cell adhesion (Carstens *et al*. 1996; Jones *et al*. 2019; Li and Burridge 2019) and growth factor-mediated signalling (Golias *et al*. 2004; Jones and Kazlauskas 2001; O’Keefe and Pledger 1983; Rudland and Jimenez de Asua 1979; Schwartz 1997). The possible impact adhesion growth factor crosstalk, seen to regulate anchorage-dependent signalling, has on AURK activation and function will have implications for multiple cellular pathways and processes. This study reveals AURKA and AURKB activity to drop on the loss of adhesion, AURKB activation (but not AURKA) recovering on re-adhesion in serum-deprived conditions. This confirms AURKB to be regulated downstream of integrin mediated adhesion in mouse fibroblasts. This regulation is independent of the cell cycle, as serum-deprived fibroblasts showed no significant change in their cell cycle profile when suspended or re-adherent.

Knowing the effect adhesion growth factor crosstalk could have on cellular signaling, we tested and find the presence of serum did increase the percentage of S-phase cells, though the loss of adhesion and re-adhesion did not affect their cell cycle profile. In cells grown in the presence of serum re-adhesion mediated activation of AURKB is blocked. This when considered with the fact that basal activation of AURKB in suspended cells is unaffected by serum, suggests the serum-mediated regulation of AURKB to be unique to its crosstalk with integrin-mediated adhesion. The fact that a short 15min serum treatment can block the recovery of AURKB in serum-deprived re-adherent cells suggests this regulation to be independent of the cell cycle and mediated by a serum component. Heat inactivation of serum (56°C, 30mins) did not affect this regulation suggesting this component to be heat resistant. This could hence be a growth factor (like EGF, FGF, TGF) or small molecules like amino acids, sugars, lipids, or hormones (Honn *et al*. 1975). The dialysis or charcoal treatment of serum (Stoikos *et al*. 2008) could further help identify the serum fraction that carries this component.

Serum is seen to regulate Aurora Kinases, driving AURKA activation at the basal body of the cell cilium in non-cycling G0/G1-phase mammalian cells, causing AURKA-dependent ciliary resorption (Pugacheva *et al*. 2007). Calcium-Calmodulin signalling is also seen to regulate the activation of AURKA (Plotnikova *et al*. 2010) and AURKB (Mallampalli *et al*. 2013), which downstream of integrins (Balasubramanian *et al*. 2007; Kwon *et al*. 2000; Shankar *et al*. 1993) through focal adhesion signalling (Giannone *et al*. 2004; Kirchhofer *et al*. 1991; Naik and Naik 2003) could regulate AURKB. Cell-matrix adhesion could also regulate the spatial localization and activation of AURKB. Its localization at membrane protrusions could be mediated by formin (FHOD1) (Floyd *et al*. 2013) which is the major formin in mouse fibroblasts and seen to be active at sites of integrin engagement in cells (Iskratsch *et al*. 2013). Adhesion is also seen to regulate membrane raft trafficking which through resident proteins like Flotillin-1, can regulate AURKB (Gomez *et al*. 2010).

To establish the impact AURKB regulation could have on adhesion-dependent signaling in WTMEFs we tested the effect absence/presence of serum has on integrin-mediated ERK and FAK activation (Eliceiri 2001). The presence of serum promotes basal ERK activation significantly more than it affects FAK or AURKB activation. With serum regulating AURKB in re-adherent cells, ERK activation in re-adherent cells was seen to be inversely correlated to AURKB activation status. AZD1152 mediated inhibition of AURKB in serum-deprived cells confirms this crosstalk, by promoting ERK activation, that now localizes at membrane ruffles. This AURKB mediated regulation of ERK could be direct or indirect, which our ongoing studies are evaluating.

ERK, a serine/threonine kinase, is an important component of Ras-Raf-MEK-ERK signalling cascade that regulates cell proliferation, differentiation, migration, senescence, apoptosis and tumorigenesis (Chang and Karin 2001; Nishida and Gotoh 1993; Sun *et al*. 2015). The early adhesion-dependent activation of ERK is reported to specifically regulate lamellipodial protrusion in re-adherent cells. This is also seen to be associated with the detection of pERK in membrane ruffles of re-adherent cells (Mendoza *et al*. 2011). A pulsatile ERK activation is seen to be responsible for the dynamic generation of protrusions (Yang *et al*. 2018). Ras recruits the MAP kinase kinase kinase Raf, which phosphorylates and activates the MAP kinase kinase MEK, which then phosphorylates and activates ERK (McKay and Morrison 2007; Pullikuth and Catling 2007) at the plasma membrane. KSR1 (kinase suppressor of Ras 1) forms a ternary complex with B-Raf and MEK, that promotes activation of ERK and allows for its recruitment to the plasma membrane (RL Bryan 2009)(McKay *et al*. 2009; Muller *et al*. 2001).

ERK regulates cell migration through its phosphorylation of myosin light chain kinase (MLCK), focal adhesion kinase (FAK), calpain, and p21-activated kinase (PAK1). ERK (ERK1 and ERK2) is activated by its phosphorylation on Thr^202/185^ and Tyr^204/187^ (human sequences) by MAPK/ERK Kinases (MEKs) (Miningou and Blackwell 2020; Seger *et al*. 1992). Upon phosphorylation, ERK is either seen to homodimerize or heterodimerize (Khokhlatchev *et al*. 1998), which enhances ERK activity but reduces its translocation of ERK monomers to the nucleus. These phosphorylation steps (Cirit *et al*. 2010; Kholodenko 2006), regulate the dissociation of MEK from ERK. This makes ERK very sensitive to the activation status of MEK. ERK inactivation is mediated by the MAPK phosphatases (MKPs) dependent removal of the Thr or Tyr phosphorylation’s (Caunt and Keyse 2013). ERK effectors localize in the cell cytoplasm, or the nucleus (Yoon and Seger 2006), with the nuclear translocation of ERK being vital in inducing gene expression and cell cycle entry.

A regulatory crosstalk between AURKB-ERK1/2 downstream of adhesion could have implications in maintaining the fidelity of the cell cycle in anchorage-dependent cells and regulating adhesion-dependent cell spreading and/or migration. *Insilico* sequence analysis of AURKB suggests that it is could also be a possible substrate for ERK (unpublished data), as has been suggested in earlier studies (Eves *et al*. 2006). Ongoing studies in the lab are testing the same and its significance in cellular function.

During mitosis, Raf-1 is a key regulator of ERK activation (Galabova-Kovacs *et al*. 2006; Minden *et al*. 1994), through the Raf kinase inhibitory protein (RKIP) (Eves *et al*. 2006), also seen to regulate AURKB activation (Eves *et al*. 2006). ERK1/2 and AURKB are both individually involved in cell spreading (Ferreira *et al*. 2013; Fincham *et al*. 2000; Floyd *et al*. 2013; Xu *et al*. 2010) and migration (Matsubayashi *et al*. 2004; Sun *et al*. 2015; Xie and Meyskens 2013; Zhu *et al*. 2014) something their regulatory crosstalk could help fine-tune. Integrin engagement and downstream signaling pathways have been reported to be different in cells that are embedded in a 3D matrix (Davidenko *et al*. 2016; Duval *et al*. 2017; Kapalczynska *et al*. 2018; Martino *et al*. 2009). ERK1/2 activation in primary human fibroblasts grown in cell-derived 3D fibronectin was found to be significantly higher (Damianova *et al*. 2008). This combined with the fact that cell cycle progression is altered in cells embedded in 3D matrix (Desmaison *et al*. 2018; Moriarty and Stroka 2018) suggests that AURKB activity might be regulated differentially in 2D vs 3D microenvironments. Evaluating the AURKB-ERK1/2 crosstalk in 3D microenvironments could also be relevant for understanding the role it could have in maintaining the fidelity of the cell cycle in anchorage-dependent cells.

In anchorage-independent cancers, AURKB and ERK1/2 have been reported to synergistically enhance tumorigenic potential and aid in radio-resistance (Marampon *et al*. 2014; Niermann *et al*. 2011). In melanoma cells, BRAF/ERK axis has been shown to control AURKB expression at the transcriptional level (Bonet *et al*. 2012). In gynaecological cancers, MEK/ERK cascade has been shown to regulate AURKB signaling to sustain colony-forming potential, invasion, and migration along with altering their radiation response (Marampon *et al*. 2014). These reports further add to the possible role the AURKB-ERK1/2 crosstalk could have in diseases like cancer.

Much remains to be figured about the functional significance of this AURKB-ERK crosstalk in cells. This study in identifying their novel regulation downstream of adhesion and growth factors highlights a regulatory pathway that could have implications for multiple cellular processes.

## SUPPLEMENTARY FIGURE LEGENDS

**Supplementary Figure 1.**
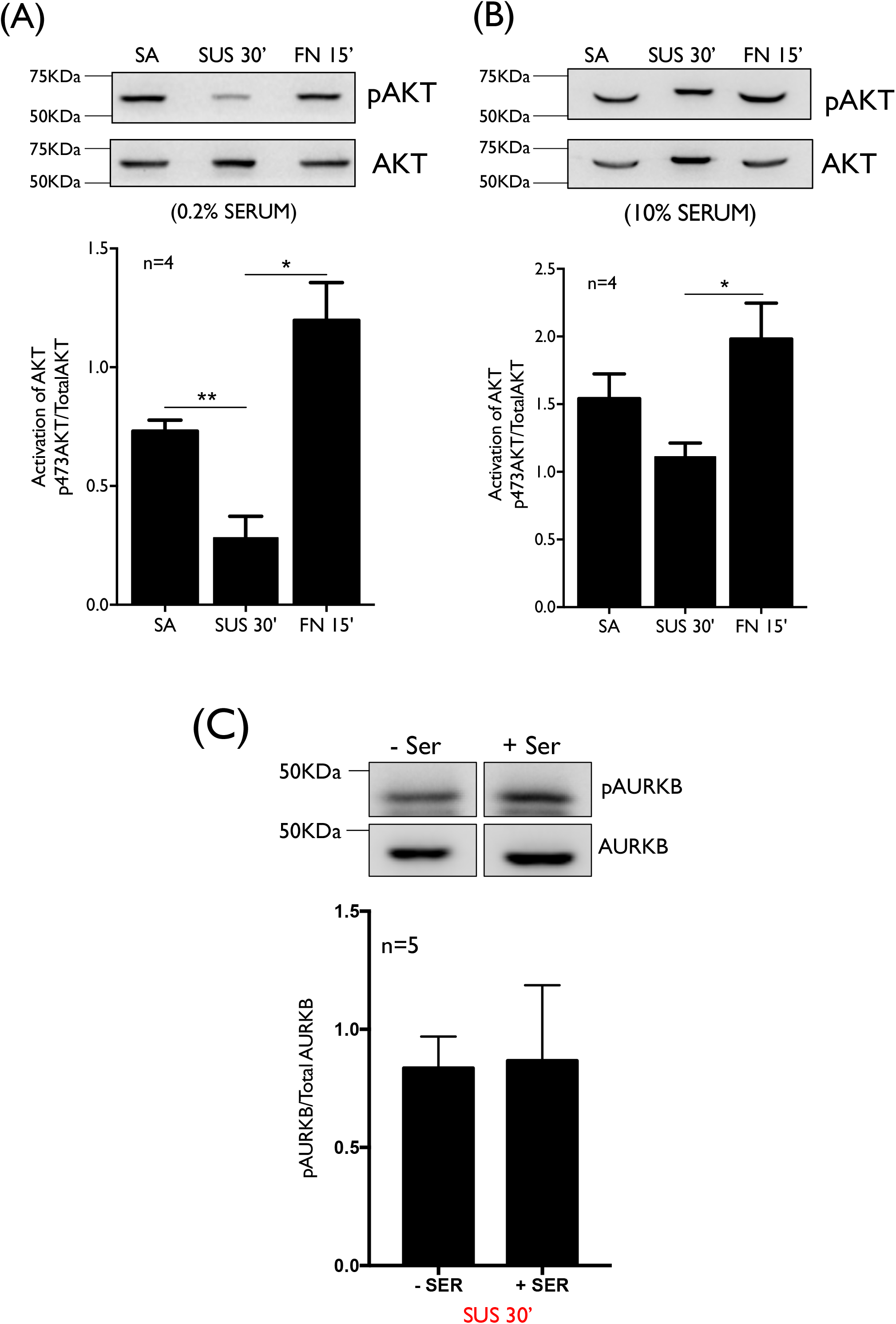
Adhesion-growth factor crosstalk dependent regulation of AKT. Western blot detection (upper panel) and quantitation (lower panel) of (A and B) phosphorylation on Tyrosine 473 residue of AKT (pAKT), total AKT in the lysates from WT-MEFs stable adherent (SA), suspended for 30 mins (SUS 30’) and re-adherent on fibronectin for 15mins (FN 15’) in presence of 0.2% (A) and 10% (B) FBS DMEM and (C) phosphorylation on the Threonine 232 residues of AURKB (pAURKB), total AURKB in lysates from WT-MEFs suspended for 30mins. The ratios of pAKT/AKT and pAURKB/AURKB are represented in the graph as mean ± SE from at least three independent experiments. Statistical analysis of all the above data was done using a two-tailed paired Student’s *t*-test and significance if any was represented in the graph (* p-value <0.05, ** p-value <0.01).

**Supplementary Figure 2.**
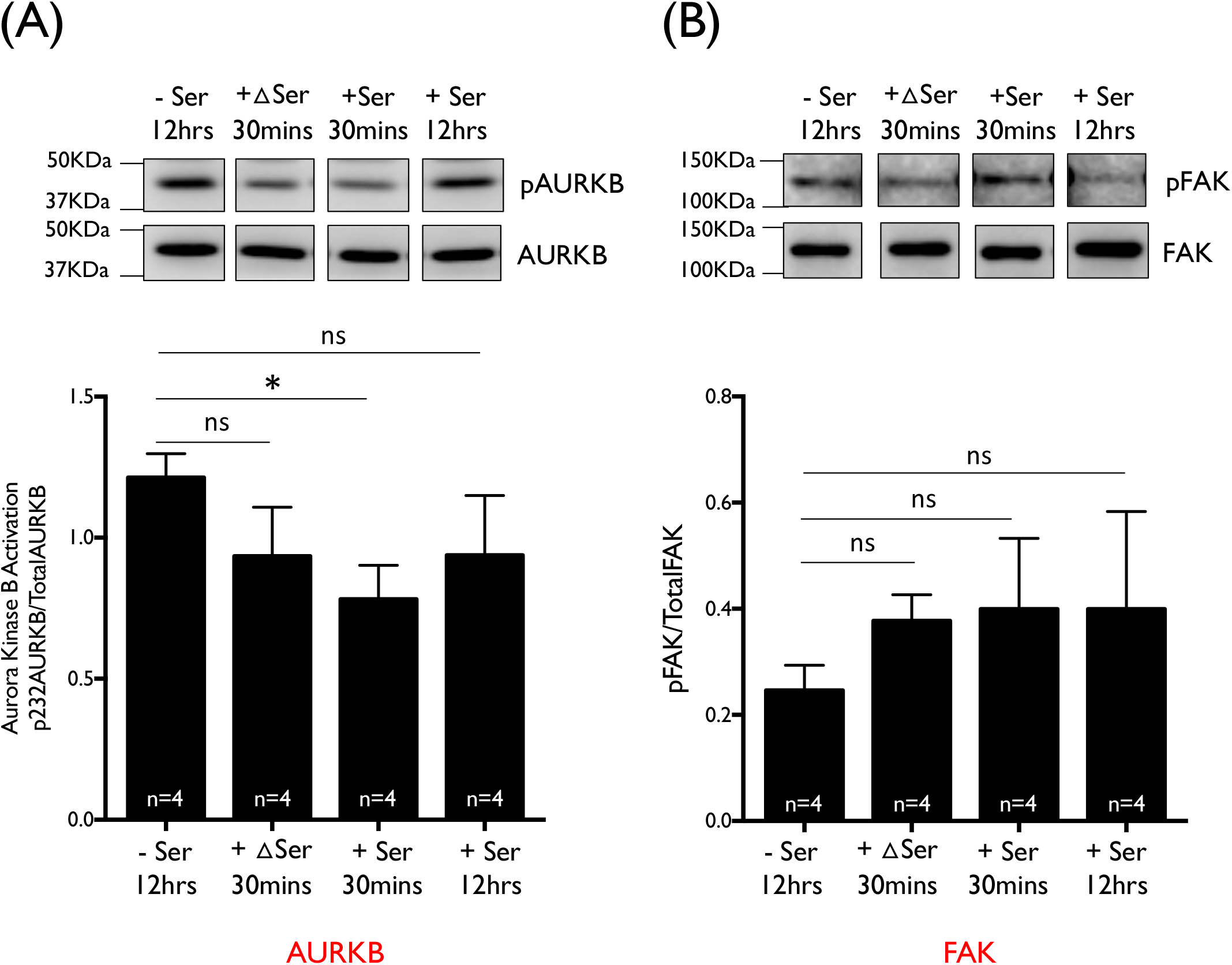
Effect of serum stimulation on AURKB activity in suspended WT-MEFs. Western blot detection (upper panel) and quantitation (lower panel) of (A) phosphorylation on the Threonine 232 residues of AURKB (pAURKB), total AURKB and, (B) phosphorylation on Tyrosine 397 residues of FAK (pFAK) and total FAK in the lysates from: serum starved WT-MEFs (-Ser 12hrs) suspended for 45mins in presence of 0.2% serum, serum starved WT-MEFs suspended for 30 mins followed by 15mins in heat inactivated 10% FBS DMEM (+ΔSer 30mins), serum starved WT-MEFs suspended for 30 mins followed by 15mins in 10% FBS DMEM (+Ser 30mins) or WT-MEFS suspended in presence of 10% FBS DMEM. The ratios of pAURKB/AURKB and pFAK/FAK are represented in the graph as mean ± SE from four independent experiments. Statistical analysis of all the above data was done using the two-tailed unpaired Student’s t-test and significance if any was represented in graph (* p-value <0.05).

**Supplementary Figure 3.**
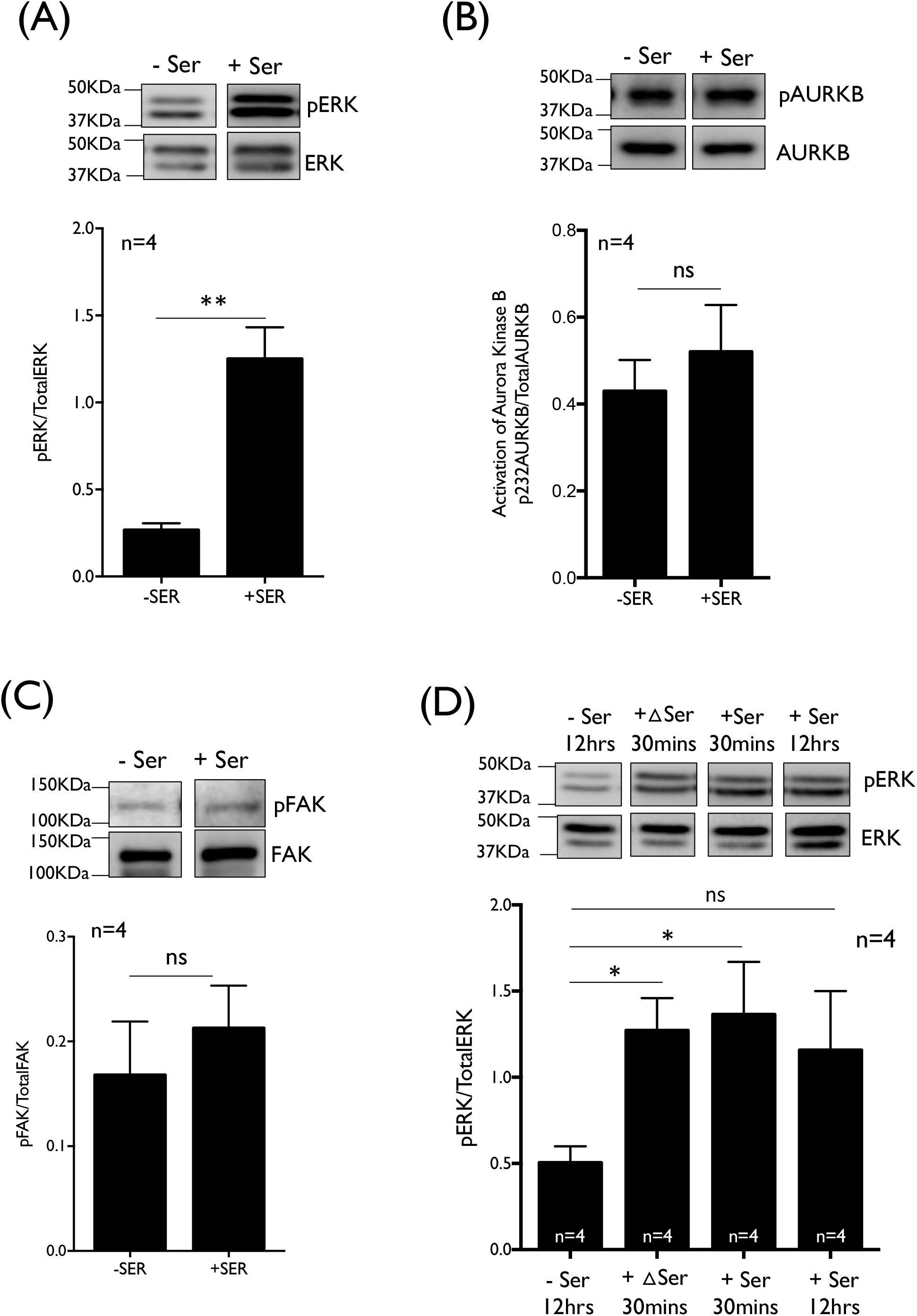
Adhesion-growth factor crosstalk dependent regulation of ERK, AURKB and FAK activity in suspended WT-MEFs. Western blot detection (upper panel) and quantitation (lower panel) of (A) phosphorylation on Threonine202/Tyrosine204 residues of p44/p42 ERK1/2 (pERK) and Total p44/p42 ERK1/2 (TotalERK), (B) phosphorylation on the Threonine 232 residues of AURKB (pAURKB), total AURKB and, (C) phosphorylation on Tyrosine 397 residues of FAK (pFAK) and total FAK in the lysates from: in the lysates from WT-MEFs suspended in absence and presence of serum. (D) Western blot detection (upper panel) and quantitation (lower panel) of phosphorylation on Threonine202/Tyrosine204 residues of p44/p42 ERK1/2 (pERK) and Total p44/p42 ERK1/2 (TotalERK) in lysates from serum-starved WT-MEFs (-Ser 12hrs) suspended for 45mins, serum-starved WT-MEFs suspended for 30 mins followed by 15mins in heat-inactivated 10% FBS DMEM (+ΔSer 30mins), serum-starved WT-MEFs suspended for 30 mins followed by 15mins in 10% FBS DMEM (+Ser 30mins) and WT-MEFS suspended in presence of 10% FBS DMEM. The ratios of pERK/ERK, pAURKB/AURKB and pFAK/FAK are represented in the graph as mean ± SE from four independent experiments. Statistical analysis of all the above data was done using the two-tailed unpaired Student’s t-test and significance if any was represented in graph (* p-value <0.05, ** p-value < 0.01).

**Supplementary Figure 4.**
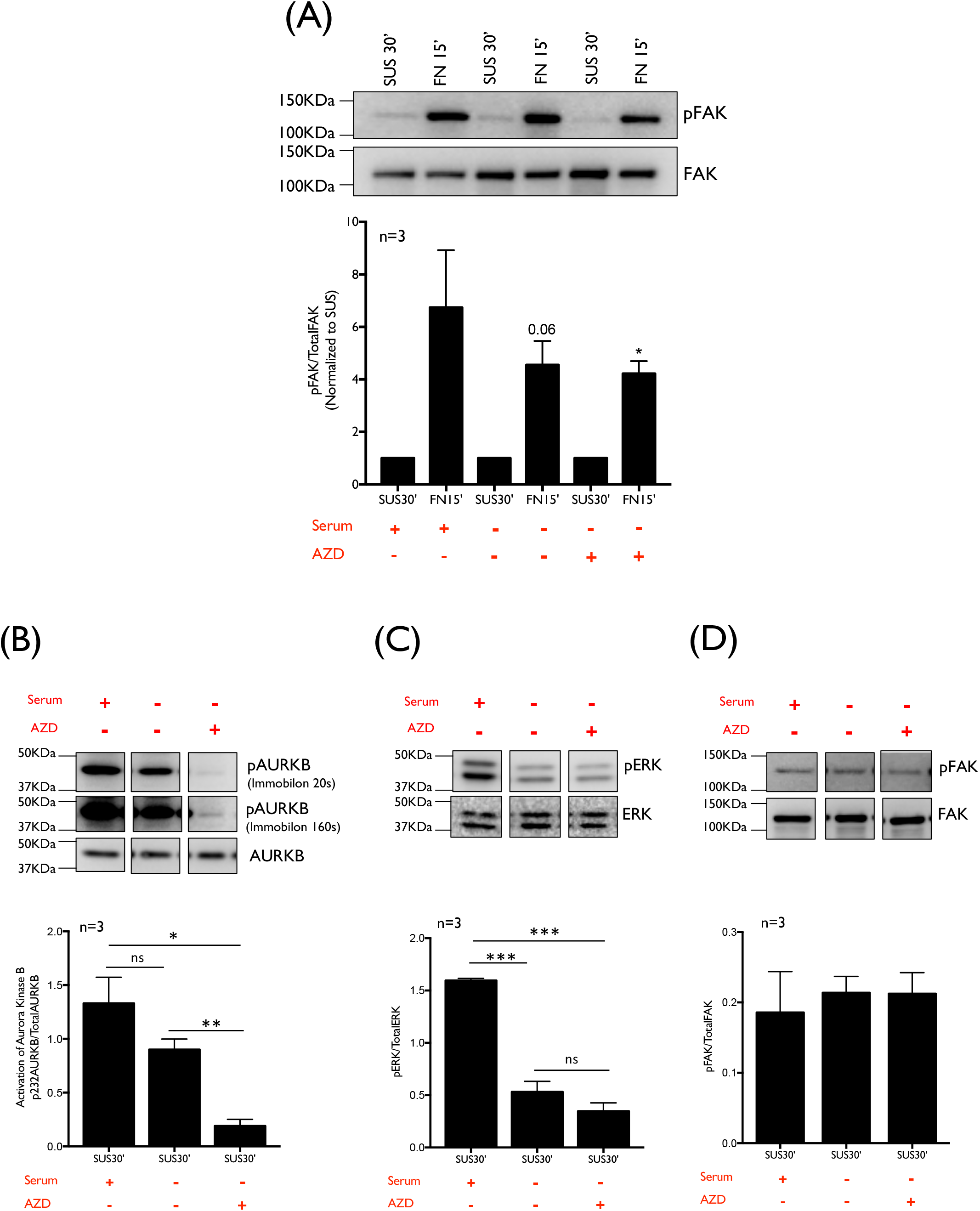
Effect of AURKB inhibition on AURKB, ERK and FAK activity in suspended WT-MEFs. Western blot detection (upper panel) and quantitation (lower panel) of (A) phosphorylation on Tyrosine 397 residues of FAK (pFAK) and total FAK in the lysates from serum starved WT-MEFs suspended for 30mins (with or without 2μM AZD1152) and re-plated on fibronectin in presence of 0.2% serum (with or without 2μM AZD1152) and WT-MEFS suspended and re-plated in presence of 10% FBS DMEM, and, (B) phosphorylation on the Threonine 232 residues of AURKB (pAURKB), total AURKB, (C) phosphorylation on Threonine202/Tyrosine204 residues of p44/p42 ERK1/2 (pERK) and Total p44/p42 ERK1/2 (TotalERK), (D) phosphorylation on Tyrosine 397 residues of FAK (pFAK) and total FAK in the lysates from 30 mins suspended cells in presence of 0.2% serum (with or without 2μM AZD1152) or with 10% serum. The ratios of pFAK/TotalFAK, pAURKB/AIRKB and pERK/ERK are represented in the graph as mean ± SE from three independent experiments and normalized to SUS (equated to 1) in case of A. Statistical analysis of all the above data was done using the single sample t-test for normalized data and two-tailed unpaired Student’s t-test for non-normalized data and significance if any was represented in graph (* p-value <0.05, ** p-value<0.01)

## Notes

### Competing Interest Statement

The authors have declared no competing interest.

